# FiberSim: a flexible open-source model of myofilament-level contraction

**DOI:** 10.1101/2021.06.11.448126

**Authors:** S. Kosta, D. Colli, Q. Ye, K. S. Campbell

## Abstract

FiberSim is a flexible open-source model of myofilament-level contraction. The code uses a spatially explicit technique, meaning that it tracks the position and status of each contractile molecule within the lattice framework. This allows the model to simulate some of the mechanical effects modulated by myosin binding protein-C, as well as dose-dependence of myotropes and the effects of varying isoform expression levels. This paper provides a short introduction to FiberSim and presents simulations of tension-pCa curves with and without regulation of thick filament transitions by myosin-binding protein C. The software was designed to be flexible (the user can define their own model and/or protocol) and computationally efficient (simulations can be performed on a regular laptop). We hope that other investigators will use FiberSim to explore myofilament level mechanisms and to accelerate research focusing on the contractile properties of sarcomeres.

**Statement of significance:** Myotropes, such as omecamtiv mecarbil and mavacamten, are new therapeutics that bind directly to sarcomeric proteins. Their clinical development has reenergized interest in mechanistic understanding of sarcomere level function. FiberSim is an open-source spatially-explicit computer model that simulates myofilament level mechanics and can predict how modulating the function of a sarcomeric protein will impact contractility. The software was specifically designed to be flexible and user-friendly and may help to further accelerate myofilament-research.

## Introduction

Different approaches exist to model myofilament-level contraction. For example, Andrew Huxley’s original work [1] simulated myofilament dynamics using a system of partial differential equations that describe the time-dependent evolution of distributions of attached myosin heads. An extension of Huxley’s approach, termed MyoSim [2] has been developed by our lab and can reproduce many features of contractile dynamics [3,4]. However, models based on Huxley’s approach do not incorporate the three-dimensional configuration of the sarcomere lattice, and do not implement all of the spatial constraints on protein interactions.

Spatially explicit models of myofilaments track of the position and status of each contractile protein in the sarcomere lattice. These models typically run more slowly than Huxley-type systems and are more complicated to program. However, they may provide a more accurate description of contractile function, as the spatial arrangement of proteins within the sarcomere is likely to impact contractile function [5].

To the authors’ knowledge, Daniel, Trible, and Chase [6] developed the first spatially explicit model of a half-sarcomere. This initial model introduced the concept of compliant realignment of myosin heads and actin binding sites and spurred further work [7], including Fenwick et al.’s study of the spatial distribution of thin filament activation [8]. Arguably the most sophisticated model to date is MUSICO, developed by Mijailovich et al. [9]. This important model simulates the activation of both thin and thick filaments and has provided important new insights into sarcomere-level function [10].

FiberSim (https://campbell-muscle-lab.github.io/FiberSim/) builds on the pioneering work described above [6–10] but was developed with special emphasis on usability and flexibility. The user can vary the biophysical properties of each contractile protein and simulate different types of protocol. FiberSim takes ∽3 minutes to simulate an isometric twitch contraction on a standard Windows laptop. This makes FiberSim more accessible to most users than prior spatially explicit models. For example, MUSICO takes ∽10 hours on a system with 192 processors to simulate a comparable experiment [10].

The following sections provide additional details about the FiberSim model and present example simulations demonstrating how myosin-binding protein C (MyBP-C) can modulate contractile function via localized effects within the sarcomere lattice.

## Methods

FiberSim describes the contractile properties of a half-sarcomere composed of thick and thin filaments. The underlying framework is similar to that described by groups led by Daniel, Chase, Tanner, and Mijailovich [6,7,9]. The code tracks the position and status of each myosin head, each binding site on actin, and each MyBP-C molecule. The software is separated into two components:

- FiberCpp is the “core model” and implements the calculations underlying the myofilament system. It is written in C++ and designed solely to run quickly and efficiently. The results of the simulations are written to files using standard formats for portability and efficiency.
- FiberPy is the “interface”. This part of the software is written in Python to facilitate rapid development and flexibility. It provides options to run simulations, analyze simulation outputs, visualize the FiberSim framework, make figures and fit simulations to experimental data.

### Computational structure

As shown in Fig. 1, FiberSim requires three input files to run a simulation:

**Fig. 1:**
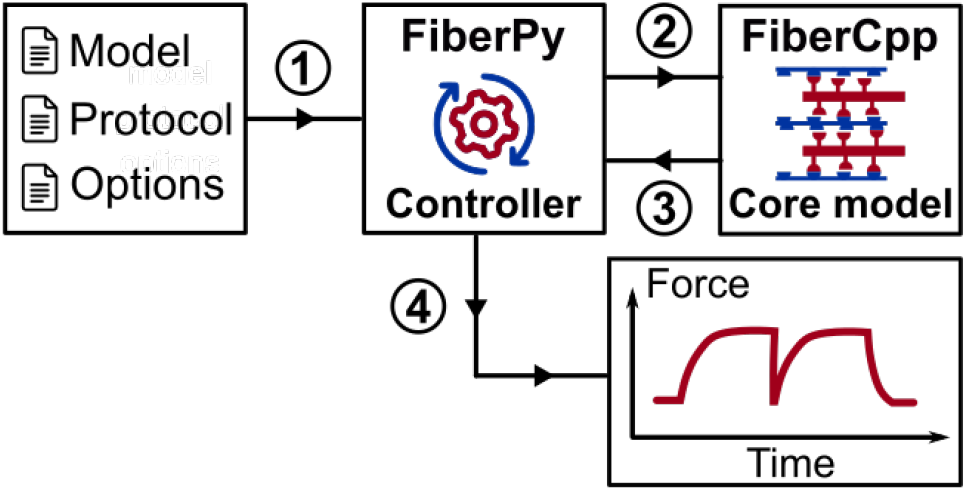
The FiberSim framework. FiberPy is the Python code that handles communication with the core model. FiberCpp is the C++ code that simulates the myofilament level biophysics. Once the simulation is complete, FiberCpp writes the results of the calculations to an output file. FiberPy analyzes the results, summarizes the data, and plots figures.

1. The protocol file is a .txt file with four columns: the time-step, the pCa value (pCa = - log_10_ [Ca^2+^]) the imposed length change (held at 0 for isometric), and the mode of contraction (length-control or force-control mode). Each new line of the protocol file represents an additional time-step.
2. The model file is a JSON file that describes the half-sarcomere lattice structure and the filament kinetics (see below). JSON is a file format for storing and transporting data that is simple and human-readable. Each parameter value comes with a “key” (see Supplementary Material) which simplifies the process of adding new features to FiberSim as understanding of sarcomere biology evolves.
3. The option file is a JSON file that specifies the computational options, such as the tolerance limit for the force-balance calculations, and whether to create log files that track the status of the filaments throughout the simulations.

### Sarcomere lattice and filament structure

A visualization of the half-sarcomere lattice in shown in Fig. 2. Myosin heads are arranged in dimers and can attach to available binding sites on the nearest thin filaments. Thick and thin filaments located at the edge of the lattice are “mirrored” on the opposite side to minimize edge effects [7,8].

**Fig. 2:**
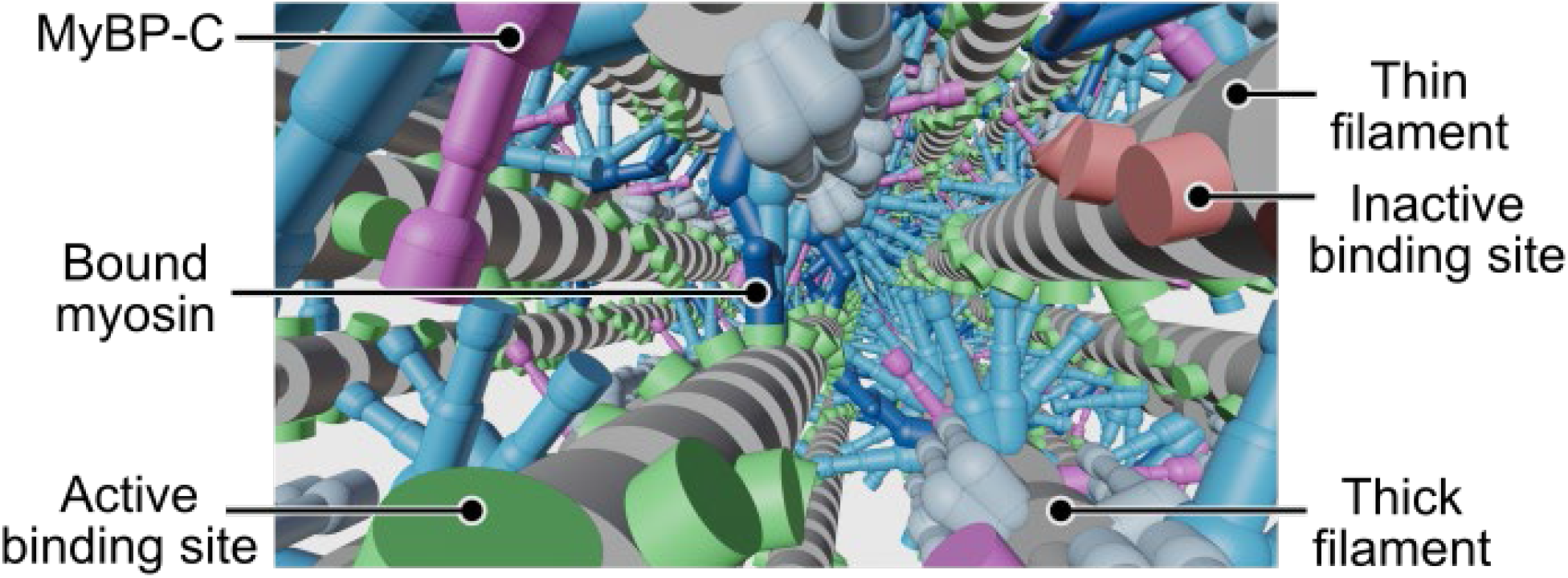
Visualization of the half-sarcomere lattice. Myosin heads from the thick filament can attach to binding sites on the surrounding thin filaments. This sarcomere lattice snapshot is generated using Blender (an open-source visualization software available at https://www.blender.org/). Blender movies showing a half-sarcomere contraction are available on the FiberSim website.

FiberSim tracks the location of filament “nodes” where either actin binding sites, myosin heads, or MyBP-C molecules are located. Fig. 3 shows a simplified 2D representation of a single half-thin and a single half-thick filament. Nodes on each filament are joined by linear springs to account for the filaments’ compliance. The thick filament is connected to the M-line by a rigid link to represent the bare zone. In 3D, titin springs link each thick filament to the six nearest thin filaments.

**Fig. 3:**
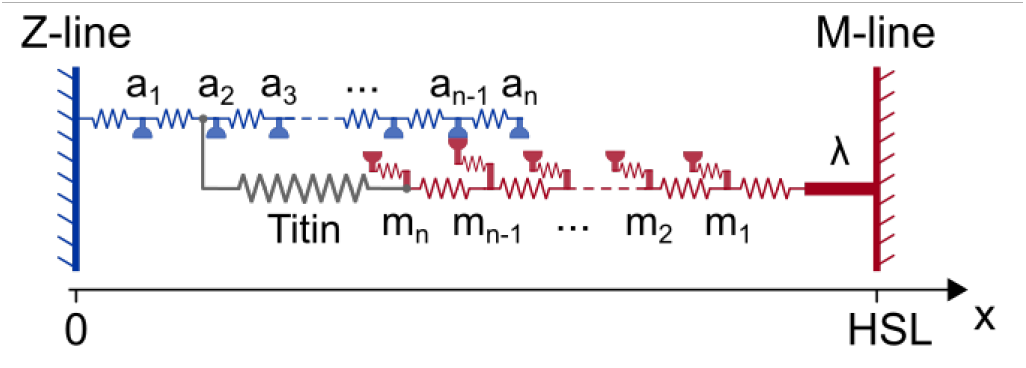
2D representation of a thin and a thick filament from the half-sarcomere lattice. Each filament is compliant, and represented as a series of nodes connected by linear springs. Actin binding sites are located on the thin filament nodes. Myosin heads and MyPB-C are located on the thick filament nodes. Myosin heads are represented as extensible springs and are able to attach to available actin binding sites. Titin molecules are represented as springs connecting the end of a thick filament to the to the six nearest thin filaments. The thick filament is linked to the M-line by a rigid link of length λ to represent the bare zone. HSL is the half-sarcomere length.

The JSON model file defines the number of filaments in the simulation as well as structural and biophysical properties including the number of binding sites per filament, the stiffness of the compliant link, and the rate functions defining binding site and myosin kinetics. An example file is provided in the Supplementary Material. This model is intended to mimic striated muscle structure so that:

- Each thin filament node holds two diametrically opposed binding sites, each of which can be available (active) or not available (inactive) for myosin binding.
- Each thick filament node holds six myosin heads arranged in three dimer pairs. Each myosin head can bind to a nearby active actin binding site.

At each simulation time-step, the x-position of every node in the lattice framework is calculated by assuming that forces are balanced at each node [6]. FiberSim performs this calculation using an iterative approach based on the conjugate gradient method [11]. This approach takes advantage of the sparsity of the stiffness matrix originally described by Daniel et al. [6]. This is much faster and requires less memory than solving the full matrix equation using direct methods.

### Thin and thick filament kinetics

Seven consecutive active binding sites form a regulatory unit, and each of the seven binding sites can simultaneously switch between an inactive or active state, depending on the calcium concentration and on the transition rate constants k_on_ and k_off_. Those parameters are defined by the user. A cooperative mechanism is also implemented and controlled by the k_coop_ parameter such that:

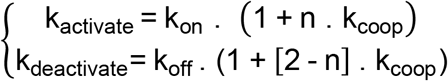

where n is the number of adjacent regulatory units in the active state (n = 0, 1 or 2).

Individual myosin heads kinetics can be defined by the user as well. In Fig. 4, a three-state myosin model is presented. The rate functions defining the probability of completing a transition within a time-step are set by the user. For each-time step, we follow the approach described in [12] to determine if a transition occurs or not.

**Figure 4:**
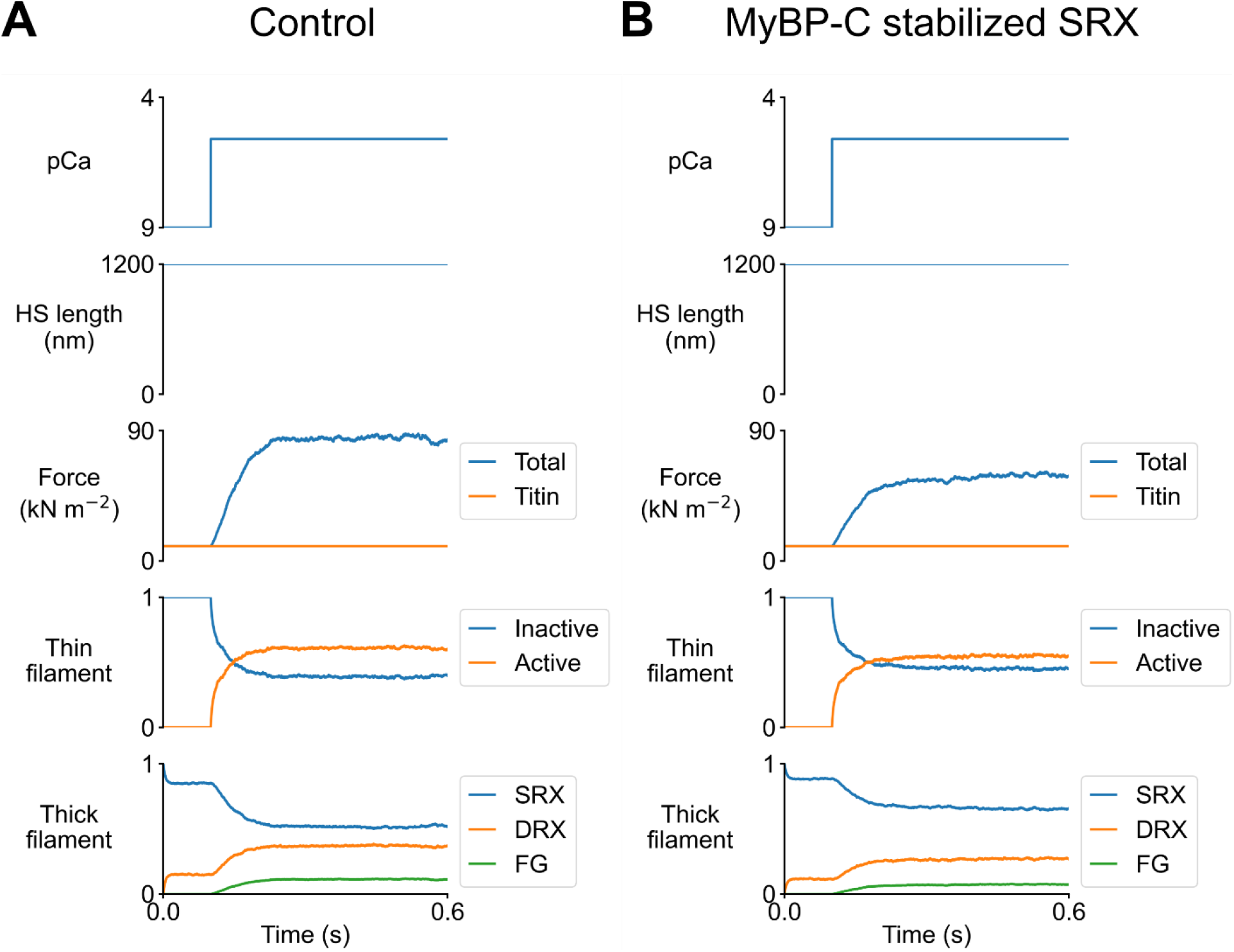
Raw data for tension-pCa curves generation. Time-traces for the different variables in the base (left) or the MyBP-C stabilized SRX (right) case at pCa = 5.6.

### Software availability

FiberSim is available for free download under the GPL3 license. Full documentation and examples are available at the project home page, https://campbell-muscle-lab.github.io/FiberSim/. Source code, bug reports, and development plans are provided at https://github.com/campbell-muscle-lab/FiberSim/. The latest release, version 1.0, is archived at https://zenodo.org/badge/latestdoi/248878056.

## Results

The results presented in this section are reproduced from examples posted at https://campbell-muscle-lab.github.io/FiberSim/pages/manuscripts/2021a/2021a.html.The simulations investigate how one potential action of MyBP-C impacts contractile dynamics. Specifically, the calculations predict how contractile force is changed if MyBP-C stabilizes the super-relaxed (SRX) state of myosin [13,14]. The authors note that MyBP-C may also impact contraction via other modes [15].

The calculations simulate a half-sarcomere lattice containing 100 thick and 200 thin filaments. Myosin heads cycle through a three-state model [12] (see Fig. 4 and Supplementary Material) that includes a super-relaxed (SRX) state. Dimers of myosin heads can transition together from the super-relaxed state (grey structures in Fig. 2) to the disordered-relaxed state (DRX, light blue structures in Fig. 2). Then each myosin head is able to bind to an available actin binding site and transition to a force-generating state (FG, dark blue structures in Fig. 2). The presence of MyBP-C (pink structures in Fig. 2) is assumed to stabilize the SRX, so that k_1_ for myosin dimers in the C-zone of the sarcomere is 70% of that for dimers in other regions of the sarcomere.

Fig. 5 show the time traces for an isometric activation protocol at pCa = 5.6 in the base case, and in the MyBP-C stabilized SRX case. Fig. 6 shows tension-pCa curves generated by running the isometric activation protocols with different pCa values. Each tension-pCa curve simulation runs in approximately 15 minutes on a regular laptop. Interestingly, there is a rightward shift of the tension-pCa curve when MyBP-C comes into play (pCa_50_ = 5.63 compared to pCa_50_ = 5.69 in the base case).

**Fig. 5:**
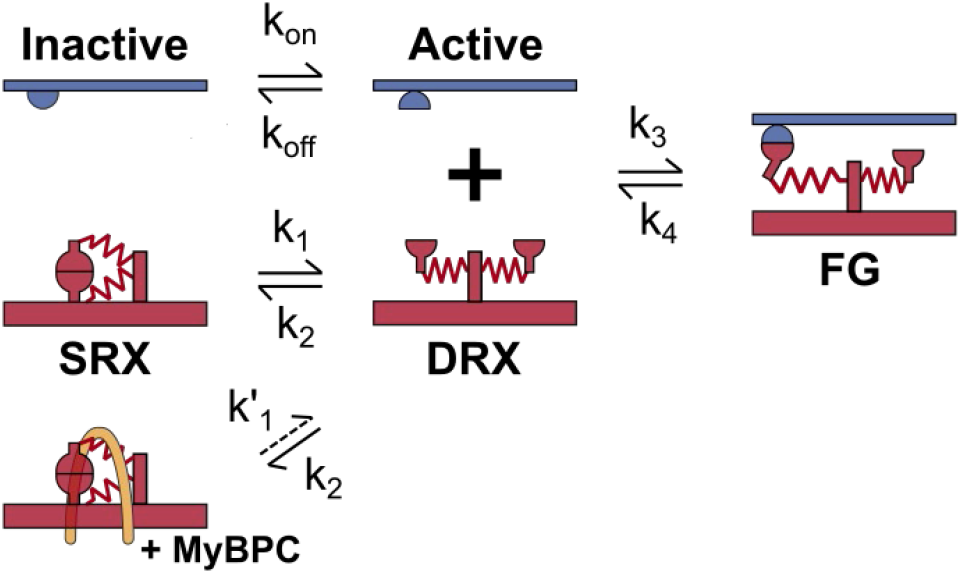
Example of a three-state myosin kinetic model. Myosin dimers (red) can transition together between the super-relaxed state and the disordered-relaxed state. Then they can transition independently to a force-generating state. Attachment to actin (blue) and force generation are only allowed if the binding site is available for binding (active state). Rate constants (k_i_) can be constant or depend on stretch, node force, or on the MyBP-C status.

**Fig. 6:**
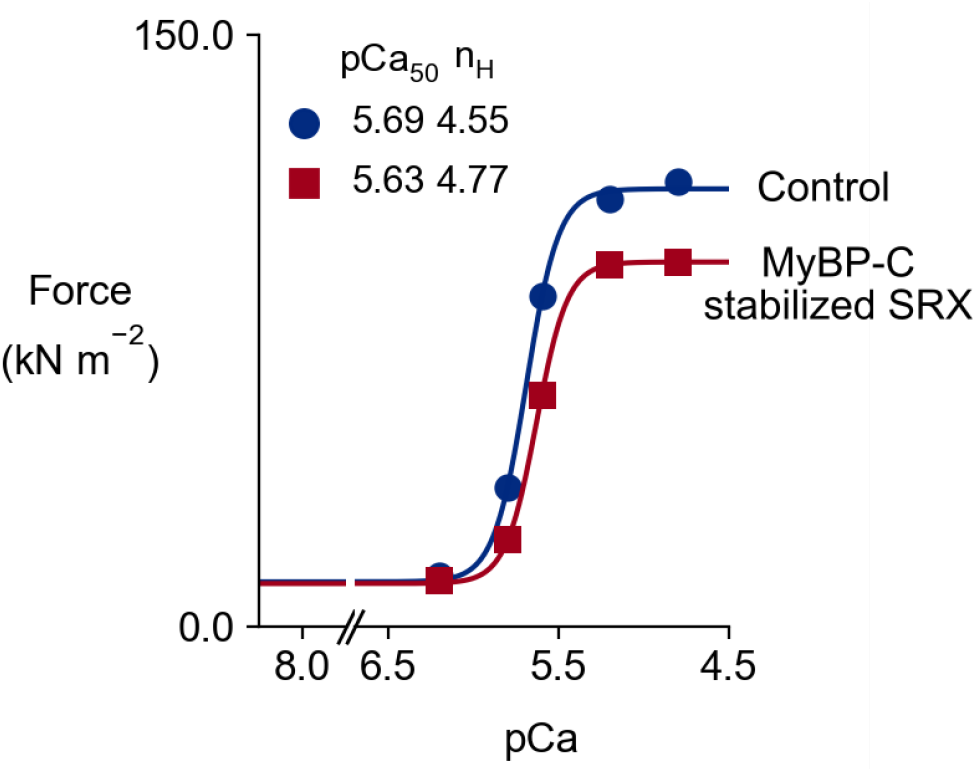
Tension-pCa curves obtained in the base case (blue), and in the MyBP-C stabilized SRX case (red).

**Figure 6:**
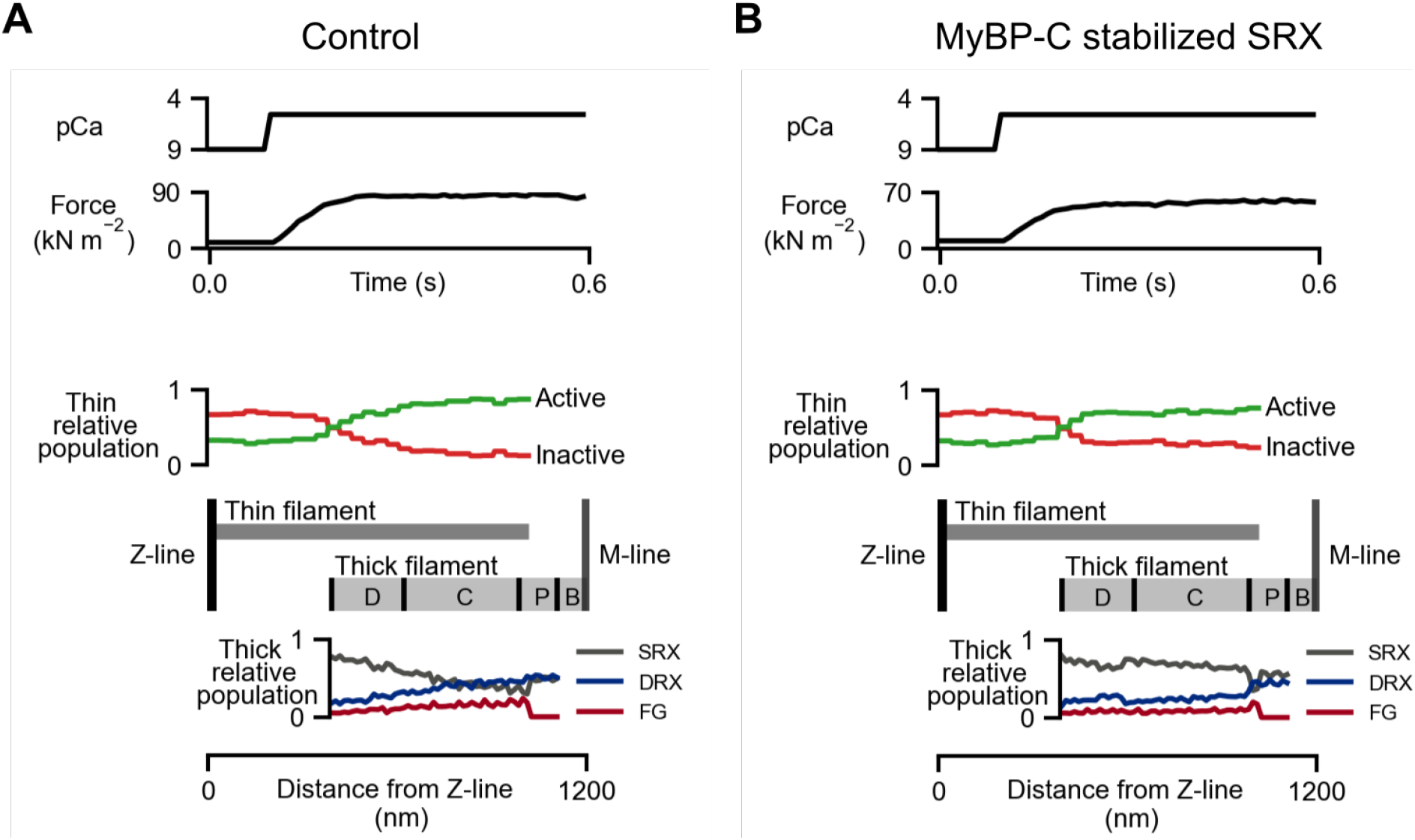
Average actin and myosin states distribution along the half-sarcomere at pCa 5.6 where isometric force is ∽70% of maximum. Left: Average distributions in the base case. Right: Average distributions with MyBP-C stabilized SRX. There is a decrease in the DRX and FG populations in the C-zone where MyBP-C stabilizes the SRX state.

Since the model is spatially explicit, the average spatial distribution of actin and myosin populations at a fixed time-step can be analyzed. In Fig. 7, those populations are plotted as a function of the distance to the Z-line. A thin and thick filament are also shown to help visualize the overlap region. In the presence of MyBP-C there is a local decrease in the average distribution of FG myosin heads in the C-zone of the thick filament, where MyBP-C decreases the recruitment rate from the SRX state. This induces a loss of calcium-sensitivity (reduction in pCa_50_) and a decrease in maximal force at full calcium activation. Movies showing the time-evolution of the spatial population distribution with and without active MyBP-C are available at https://campbell-muscle-lab.github.io/FiberSim/pages/manuscripts/2021a/2021a.html.

## Discussion

FiberSim runs on standard PCs and can simulate typical experiments in a few minutes. The software was designed to be flexible and users can define their own simulation protocols, adjust the structural details of the filament models and implement different types of myosin and MyBP-C kinetics. More demos, including isometric twitch contractions and simulations of force-velocity relationships) are provided on the FiberSim website.

The spatially-explicit aspect of FiberSim allows for precise description of local behaviors at the molecular level. For instance, myosin heads can only bind to nearby sites. This feature is not incorporated in Huxley-type models. Similarly, SRX to DRX transitions can depend on the local force in the thick filament. This leads to intra-half-sarcomere effects such as in Fig 7 where the DRX population increases towards the M-line as the force in the thick filament backbone accumulates. MyBP-C also exhibits local effects due to its restricted location within the sarcomere. While some of these effects can be mimicked using complex sets of differential equations [12], they arise naturally in FiberSim because of its underlying architecture.

## Supporting information

model file for control simulations

model file for simulation with MyBP-C stabilized SRX simulations

## Author contributions

SL wrote the manuscript, helped develop the website and demonstrations, and optimized and tested FiberCpp. DC worked on prototypes of the code-base and developed the website and GitHub repository. QY helped optimize the force-balance algorithm in the core model. KC planned the overall project, wrote the first versions of FiberCpp and FiberPy, and helped develop the demonstrations.

## Funding

Supported by NIH HL146676, HL148785, TR0001998, and AHA TP135689 to KSC and AHA 929744 to SK.

## Notes

https://campbell-muscle-lab.github.io/FiberSim/

### Competing Interest Statement

The authors have declared no competing interest.

